# Insights to HIV-1 coreceptor usage by estimating HLA adaptation with Bayesian generalized linear mixed models

**DOI:** 10.1101/2022.07.06.498925

**Authors:** Anna Hake, Anja Germann, Corena de Beer, Alexander Thielen, Martin Däumer, Wolfgang Preiser, Hagen von Briesen, Nico Pfeifer

## Abstract

The mechanisms triggering the human immunodeficiency virus type I (HIV-1) to switch the coreceptor usage from CCR5 to CXCR4 during the course of infection are not entirely understood. While low CD4^+^ T cell counts are associated with CXCR4 usage, a predominance of CXCR4 usage with still high CD4^+^ T cell counts remains puzzling. Here, we explore the hypothesis that viral adaptation to the human leukocyte antigen (HLA) complex, especially to the HLA class II alleles, contributes to the coreceptor switch. To this end, we sequence the viral *gag* and *env* protein with corresponding HLA class I and II alleles of a new cohort of 312 treatment-naive, subtype C, chronically-infected HIV-1 patients from South Africa. To estimate HLA adaptation, we develop a novel computational approach using Bayesian generalized linear mixed models (GLMMs). Our model allows to consider the entire HLA repertoire without restricting the model to pre-learned HLA-polymorphisms as well as to correct for phylogenetic relatedness of the viruses within the model itself to account for founder effects. Using our model, we observe that CXCR4-using variants are more adapted than CCR5-using variants (p-value =1.34e-2). Additionally, adapted CCR5-using variants have a significantly lower predicted false positive rate (FPR) by the geno2pheno[coreceptor] tool compared to the non-adapted CCR5-using variants (p-value =2.21e-2), where a low FPR is associated with CXCR4 usage. Consequently, estimating HLA adaptation can be an asset in predicting not only coreceptor usage, but also an approaching coreceptor switch in CCR5-using variants. We propose the usage of Bayesian GLMMs for modeling virus-host adaptation in general.

**Author summary:** Viral control is currently our only counter mechanism against HIV-1 with no practicable cure nor a vaccine at hand. In treatment-naive patients, HLA adaptation and coreceptor usage of HIV-1 play a major role in their capability to control the virus. The interplay between both factors, however, has remained unexplored so far. Assessing the degree of viral HLA adaptation is challenging due to the exceptional genetic diversity of both the HLA complex and HIV-1. Therefore, current approaches constrain the adaptation prediction to a set of p-value selected HLA-polymorphism candidates. The selection of these candidates, however, requires extensive external large-scale population-based experiments that are not always available for the population of interest, especially not for newly emerging viruses. In this work, we present a novel computational approach using Bayesian generalized linear mixed models (GLMMs) that enables not only to predict the adaptation to the complete HLA profile of a patient, but also to handle phylogenetic-dependencies of the variants within the model directly. Using this light-weight approach for modeling (any) virus-host adaptation, we show that HLA adaptation is associated with coreceptor usage.

## Introduction

Without the prospect of a vaccine or a cure within reach, viral control is one of the major pillars for combating the HIV pandemic [1]. The coreceptor usage of HVI-1 affects the ability to control the virus. Apart from the CD4 receptor, HIV-1 needs a coreceptor for successful cell entry. Only two coreceptors have clinical relevance, CCR5 and CXCR4 [2]. Depending on their coreceptor usage, HIV-1 isolates are classified into R5-capable variants (CCR5 usage), X4-capable variants (CXCR4 usage), or R5X4-capable variants (CCR5 and CXCR4 usage) [3, 4]. While R5 variants are known to dominate early infection [5], a switch to X4 at later stage of infection occurs in roughly 50% of patients infected with subtype B HIV-1 associated with increased depletion of CD4^+^ T cells, faster progression to AIDS, and a higher mortality rate [4, 6–8]. In patients infected with subtype C HIV-1, a switch to CXCR4 usage is observed less frequently compared to subtype B [9]. Recent studies suggest that an increase in subtype C X4 variants might emerge with the increasing access to antiretroviral drug treatment and the ongoing evolution of the subtype C HIV epidemic [10].

The importance of accurate determination of coreceptor usage has increased with the approval of entry inhibitor drugs that target the CCR5 coreceptor. A determinant of coreceptor usage is the *env* protein of HIV-1. Currently, phenotypic [11–14] and genotypic [15–22] tropism assays still have difficulties accurately detecting minority populations of X4-using variants, which might lead to a predominance of X4 usage after treatment with a CCR5 antagonist. In addition, it is not only important to predict the correct coreceptor usage, but it would be of advantage to predict how close the variant is to a coreceptor switch.

Though the clinical significance of the coreceptor usage is well studied, the trigger mechanisms behind the coreceptor switch from R5 to X4 variants remain unsolved. The emergence of X4-capable variants is associated with a decrease in N-linked glycosylation of the envelope glycoprotein *env* of HIV-1 [23]. Glycosylation is a viral mechanism to mask conserved amino acids from antibody recognition, such that X4-capable variants should be more prone to antibody neutralization in theory. For antibody development, B cells have to be activated by CD4^+^ T cells. Thus, concurrent CD4^+^ T cell depletion counteracts this mechanism. How X4 variants can emerge with still high CD4^+^ T cells remains inconclusive. However, this is of great importance, since patients with intermediate to high CD4^+^ T cells contradict the current typical clinical indicators for a potential coreceptor switch such as low numbers of CD4+ T cells.

The potential interplay between HLA adaptation and coreceptor usage has not been explored so far. Viral adaptation to the immune system includes the emergence of viral escape mutations to the host’s individual HLA profile. The central role of HLA molecules is to bind peptides and present them on the cell surface to compatible T cells, which are part of the adaptive immune response. T cells are HLA-restricted, meaning that they recognize only a specific HLA-antigen complex. There are two major HLA classes — HLA class I and HLA class II. HLA class I molecules exist on all nucleated cells and bind to (self and pathogen-derived) antigens degraded from synthesized proteins in the cytosol. The corresponding HLA-antigen complex is recognized by specific CD8^+^ T cells. HLA class II molecules only occur on professional antigen-presenting cells that are able to uptake pathogens and proteins from extracellular fluid by phagocytosis or endocytosis. Thus, HLA class II molecules bind pathogen-derived antigens degraded from extracellular proteins in the vesicular compartment of the cell. The corresponding HLA:antigen complex is recognized by CD4^+^ T cells. The emergence of a mutation that hinders the successful building of the HLA-antigen complex, a so-called escape mutation, allows HIV-1 to evade a T cell-mediated immune response [24, 25].

High-throughput technologies have enabled large-scale population studies to identify many HLA-restricted polymorphisms (HLA footprints) and their role on viral control [26–31]. A prominent example is the influence of the HLA-B*27 and the HLA-B*57:01 allele on disease progression [32, 33]. Determining virus-host adaptation experimentally and computationally on an individual level is challenging due to the extraordinary genetic diversity of both the HLA complex and HIV-1. HLA adaptation models usually focus on viral polymorphisms that likely emerged due to the patient’s HLA profile. This approach requires the general consideration of the extreme large number of possible HLA alleles in the population and viral polymorphisms while modeling the fact that only few HLA alleles have a potential influence on a particular polymorphism. Current computational approaches [34, 35] tackle the complex modeling task by carrying out many rounds of preselection, including the identification of potential HLA-polymorphism candidates on large-scale cohort data and additional greedy feature selection steps to select the HLA alleles per polymorphism within the model, such that potential sites and HLA alleles might get disregarded based on significance threshold values. Since human populations and HIV subtypes display substantial genetic differences, such approaches require a large amount of data for every group of interest. Correcting for potential phylogenetic relatedness of the viral sequences used within the model as proposed by [36] is currently implemented by incorporating a transmission probability that has to be learned in a separate model. While HLA-1 restricted escape mechanisms to CTLs have been studied in detail, only few studies exist that have analyzed the impact of HLA-restricted CD4^+^ T cell escape polymorphisms [35] on viral control. In total, there is currently no available approach to estimate viral adaptation jointly to HLA class I and class II. Moreover, the available approaches require rather complex training steps to be used on new data.

In this study, we investigate the hypothesis that coreceptor usage is associated with the adaptation of the virus to the host’s HLA system, especially to the HLA class II alleles. We explore the novel possibility that viral adaptation to the HLA class II molecules would mask the virus from recognition by CD4^+^ T cells, such that no B cells are activated, and, thus, no antibodies are developed despite still high numbers of CD4^+^ T cells. Escape mutations in the rather conserved *p24* protein of HIV-1, which is involved in forming the viral capsid, emerge more likely under substantial fitness cost [37, 38]. Therefore, we estimate viral adaptation to the patient’s HLA profile only based on the *p24* protein of the *gag* gene as done previously [39–44]. This study requires a data set consisting of (1) the envelope protein sequences of the virus for determining the coreceptor usage, (2) the p24 protein for estimating the HLA adaptation, and (3) the HLA class I and II profile of the corresponding host. Chronically-infected HIV-1 patients are more likely to harbor viruses that have accumulated escape mutations to the HLA system due to the longer exposure to the human immune system. In treatment-naïve patients, the viral evolution is not restricted by selection pressure from drug exposure and is more able to mutate towards escape variants with respect to the immune system. Current available data sets often lack HLA class II allele information or have not sequenced the envelope sequence of the virus. Thus, we sequence the viral envelope gene env as well as the viral *gag* (*p24*) gene, and genotype the corresponding HLA class I (HLA-A, HLA-B, HLA-C) and II genes (HLA-DRB1, HLA-DQB1, HLA-DPB1) of the host in a new cohort of 312 treatment-naive, subtype C, chronically-infected HIV-1 patients from South Africa.

To jointly model HLA class I and class II adaptation, we develop a novel computational approach. In detail, the adaptation of a particular amino acid in a viral sequence to the host HLA profile is inferred using phylogeny-corrected, multinomial, Bayesian generalized linear mixed models (GLMMs). Without the need for an additional model, GLMMs allow to correct for phylogenetic relatedness of the variants directly by modeling the between-subject correlation as a group-level effect. Using a Bayesian setting allows to learn feature importance directly within the model by applying the horseshoe prior on all HLA class I and class II alleles of the data set and without the need for additional preselection steps or a large amount of data. The horseshoe prior is used in sparse model settings to shrink the majority of the coefficients to zero by having the point mass at zero and symmetric fat tails [45].

## Materials and methods

### Study cohort

Patients (male and female) who attended Wellness, Antenatal and HIV Clinics in the Durbanville and Stellenbosch regions of the Western Cape were recruited. Only patients older than 18 years were selected. Most of the patients were assumed to be in the chronic stage of the infection. Inclusion was based on recent diagnosis of HIV-1 infection (within the previous 6 months). In total, samples from 329 HIV-infected individuals were available. Subtype C was confirmed for 317 of the 329 samples using the COMET Tool [46]. Patients on antiretroviral were excluded from the analysis, resulting in a total of 312 patients. For each patient, clinical parameters such as sex, age, ethnicity, CD4 count, and viral load were collected. In addition, the HIV-1 genes *gag* (p24) and *env* were sequenced and the patients’ HLA I and II genes were genotyped.

### Ethical statement

PBMC and plasma samples from HIV-1 positive donors were provided by Stellenbosch University with the written informed consent of the donors. Sample collection was approved under the following ethical statement “VIROLOGICAL AND IMMUNOLOGICAL CHARACTERIZATION OF CRYOPRESERVED BLOOD AND VIRUS SAMPLES” PROJECT NUMBER: NO7/06/13

### Molecular methods

HIV status was confirmed with a serological test (Architect HIV Ab/Ab Combo, 3rd generation) on serum according to the manufacturer protocol. After the surface staining of PBMCs by incubation with a monoclonal mouse anti-human antibody coupled to fluorescent dyes, the quantification of cells expressing the CD4 antigen was measured by FACS analysis. Acquisition and analysis was performed on FACs flow cytometer using Cell Quest software.

HIV-1 deep sequencing was performed using previous described protocols [47]. Analysis of deep sequencing data was performed using an internally-developed analysis pipeline, where sequence reads in the form of FASTQ files were processed and aligned via a multi-step method.

HLA genotyping was performed using the following protocol. Genomic DNA was isolated from 200 *μ*l of EDTA-anticoagulated blood using the QIAamp DNA Blood Mini Kit (QIAGEN, Hilden, Germany). Long-range PCR primers amplified the full-length of HLA class I genes (A, B, C) from 5’-to 3’-UTR. Class II genes (DPB1, DQB1, DRB1) were amplified from exon 2 to 3’-UTR. Fragment sizes were estimated to be around 3000 bp for Class I genes and 6000 bp for Class II genes, respectively. The PCR solution contained 1 x Phusion GC buffer (including 1.5 mM MgCl2), 200 *μ*M dNTPs, 1 M Betaine, 8 *μ*g Bovine Serum Albumin (BSA), 0.4 U Phusion Hot Start II High-Fidelity DNA Polymerase (Finnzymes, Vantaa, Finland), 0.5 *μ*M of each primer and 90 ng of DNA in a total volume of 20 *μ*l. After initial denaturation at 98°C for 1 minute, 35 cycles of 98°C for 10 seconds, 65°C for 20 seconds, and 72°C for 4 minutes were performed, followed by a final extension at 72°C for 20 minutes. Agarose gel electrophoresis was used to confirm amplification and correct fragment size as well as to check for non-specific product contamination. The 3 HLA class I and class II amplicons for each individual were pooled and afterwards purified with the Agencourt AMPure XP system (Agentcourt Bioscience, Beverly, MA, United States) according to the manufacturer’s protocol to inactivate unconsumed dNTPs and to eliminate extraneous primers before library preparation. These pooled amplicons then comprised a single sample. Concentrations were measured on a FLUOstar OPTIMA microplate fluorimeter (BMP LABTECH, Ortenberg, Germany) using the Quant-iT PicoGreen assay (Invitrogen, Carlsbad, CA, United States). Sample libraries for NGS were then prepared with the Nextera XT DNA Sample Prep Kit (Illumina, San Diego, CA, United States) according to the manufacturer’s protocol, including distinct DNA fragmentation, end-polishing, and adaptor-ligation steps. Through the adaptor, every sample was finally labeled with a unique identifier sequence. Sequencing was carried out then on the Illumina MiSeq Personal Sequencer (Illumina, San Diego, CA, United States) as described by the manufacturer.

### Coreceptor prediction

Coreceptor usage is predicted using the well-established tool geno2pheno[coreceptor] [17] on the viral envelope sequences. The provided false-positive rate (FPR) corresponds to the confidence with which the sequence is classified as X4-capable. The higher the FPR, the more likely the sequence is not X4-capable, but R5. Viral strains with an FPR cutoff less than 20% are classified as X4-capable, otherwise as R5-capable according to the European Consensus Group on clinical management of HIV-1 tropism testing [48].

### Estimating HLA adaptation

Assuming independence of all sites in the viral sequence, we define the adaptation of a sequence to its host HLA profile as the adaptation of each frequent single amino acid site in the sequence to the HLA profile. Moreover, though every patient is infected by a quasispecies of viruses, we only consider the consensus sequence as in previous approaches. In order to correct for potential phylogenetic relatedness of the viral sequences used within the model as proposed by [36], we also incorporate the phylogeny of the viral sequences into the model. Thus, our model requires the amino acid sequences of the viral p24 protein, the corresponding host’s HLA I and II alleles, and the phylogeny between the viral sequences for learning the HLA adaptation (training). For each frequent site, we infer a model (HLA model) to estimate the likelihood that the site is under HLA pressure as well as a hypothetical model (baseline model) that computes the likelihood that the site is not under HLA pressure. HLA adaptation of the complete protein is then defined as a function over the product of the per-site likelihood ratios of the HLA model against the baseline model. Each per-site model is built using multinomial Bayesian generalized linear mixed models (GLMMs).

In the following, we formalize the per-site model and the final adaptation score. Afterwards, we present the selection process of the frequent sites. Since each per-site model is built using Bayesian GLMMs, we provide a brief introduction to Bayesian GLMMs and their benefit over classical GLMMs and phylogeny-corrected LMMs. In addition, we provide a section on the model specification for each per-site model.

#### Notation

Let *S* be a random variable representing the set of all possible HIV-1 amino acid sequences of a particular protein of length *L*. A particular sequence *s* is a realization of S covering all sites *l* = 1,…, *L* of the protein. A particular site *s_l_* can be realized by any amino acid (aa). Since we do not have enough power to find an HLA-restricted polymorphism at a very conserved site, we restrict the sites to *m* frequent single amino acid sites *s_j_* with *j* = 1,…, *m*, which are defined by sites that vary over the set of all HIV-1 sequences in their amino acid realization. A site is defined as frequent, if the particular amino acid is observed in at least 1% of the sequences.

The host immune system is represented by the HLA alleles of the HLA I and HLA II genes. The HLA profile of an individual consists in our case of six (homozygous in all genes) to 12 (heterozygous in all genes) different HLA alleles. Let *H* represent the set of all possible HLA I and HLA II alleles. A particular HLA profile *h* is encoded as a binary vector with zeros everywhere, apart from the positions corresponding to the HLA alleles of the HLA profile. Note, thereby homozygosity is not modeled.

We model adaptation as the conditional probability that a sequence s occurs under pressure from the host HLA profile similarly to [34]:

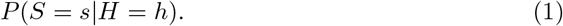

Assuming independence among sites and relevance of only frequent sites, the conditional probability over the sequence *s* can be decomposed to the product over the conditional probabilities over all *m* frequent sites *s_j_* (per-site model):

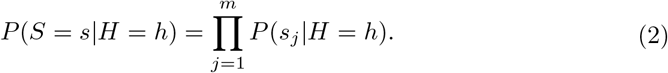

Similarly, a hypothetical model estimating the likelihood of the sequence s without any HLA pressure is defined as:

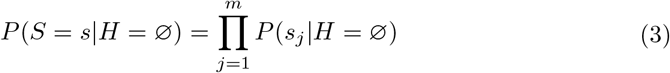

The conditional probabilities *P*(*s_j_* |*H* = *h*) and *P*(*s_j_* |*H* = ∅) for each site are referred to as the HLA model and the baseline model, respectively.

#### Identification and encoding of frequent sites

Since all patients in the study are infected with the subtype C variant of HIV-1, we align all nucleotide sequences to the subtype C consensus sequence using the alignment tool MAFFT (version 7.407) [49]. The subtype C consensus sequence is retrieved using the HIV Sequence Alignments tool from the Los Alamos HIV sequence database (www.hiv.lanl.gov/content/sequence/NEWALIGN/align.html). We correct and translate the nucleotide alignment using the Codon Align Tool from the Los Alamos HIV sequence database (www.hiv.lanl.gov/content/sequence/CodonAlign/codonalign.html). The alignment positions are mapped to the corresponding HXB2 reference gene with Genbank accession ‘AAB50258.1’ (gag) using the alignment tool MAFFT (version 7.407) [49]. Ambiguous amino acids X are not considered and set to NA. Frameshifts and stop codons are disregarded and set to gaps. Each site in the sequence *s* with at least two frequent (1% prevalence) amino acid variants is selected as potential site *s_j_* under HLA pressure. For each frequent site and each hypothesis (HLA and baseline model), a multinomial Bayesian generalized linear mixed model is built, where each frequent amino acid is considered a class, and all non-frequent amino acids are grouped together to an ‘OTHER’ class.

#### Bayesian generalized linear mixed models

We model the conditional probabilities for site adaptation (see Eq. 2 and Eq. 3) using separate multinomial Bayesian generalized linear mixed models (GLMMs). GLMMs are tailored for data with non-normal response distributions and dependency structures in the observations by combining the properties of generalized linear models (GLMs) [50, 51] and linear mixed models (LMMs). While GLMs model non-normal response distributions (such as binomial) via link functions of the means (e.g. logistic regression), LMMs enable to model not only population-level effects but also group-level effects assuming dependency structures in the samples. Mathematically, GLMMs have the following form excluding the residuals (*ϵ*) [52]:

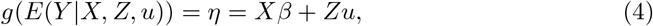

where *Y* is the response variable, *β* and *u* the coefficients for the population and group-level effects, respectively, *X* and *Z* the corresponding design matrices and *g*(*x*) a link function relating the response *Y* to the linear predictor *η*. Thus, between-subject correlations, like the phylogenetic relatedness of some viruses, can be modeled as a group-level effect.

While *y*, *X* and *Z* are given by the data, *β* and 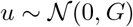 are unknown and have to be estimated. We use Markov chain Monte-Carlo (MCMC) based Bayesian GLMMs, since they are more robust and accurate in their parameter estimations of the group-level effects in contrast to classical maximum likelihood (ML) and restricted maximum likelihood (REML) methods [53]. In non-Bayesian frameworks, the group-level effect vector *u* is treated as part of the error term and thus likelihood computation requires the integration over the likelihood of all group-level effects, which might be analytically intractable for complex group-level structures [54]. In Bayesian settings where posterior distributions of the parameters are estimated by combining likelihood and prior distributions, both *u* and *β* are treated as parameters, allowing more accurate variance estimates for the group-level effects. We use the MCMC Bayesian GLMM implementation of the R [55] package brms [56] that provides an interface to the STAN software [57]. By implementing Hamiltonian Monte Carlo [58] and the No-U-Turn Sampler (NUTS) [59], Stan allows for faster convergence compared to conventional MCMC methods. Another advantage of Bayesian models is the possibility to include the prior information of the parameters into the model. The prior knowledge that only few HLA alleles have potential influence on a variant site [31, 34] is modeled using the horseshoe prior that has a global parameter *τ* shrinking most of the coefficients to zero and a local parameter λ, which is a heavy-tailed half-Cauchy (*C*^+^ (0, 1)) prior, allowing some coefficients to escape the shrinkage [45]. Thus, the the horseshoe prior for the *D* population level coefficients ***β*** = (*β*_1_,…, *β_D_*) has the following form:

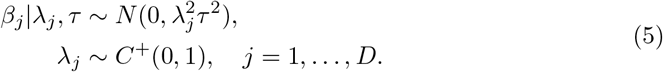

In addition, we regularize the horseshoe prior by setting the ratio of the expected number of non-zero coefficients to the expected number of zero coefficients to 10%. All other parameters of the horseshoe prior are set to default. For the remaining coefficients the default priors of the brm function are used (non or very weakly informative priors).

#### Estimating per-site adaptation

For each frequent site, we model an HLA model (see Eq. 2) and a baseline model (see Eq. 3) using multinomial Bayesian generalized linear mixed models (GLMMs) as implemented by the brms package [56] in R [55]. Both models estimate the probability distribution of each site *s_j_* spanning over the space *Y* of all frequent amino acid variants (and ‘OTHER’ for the non-frequent variants’) conditioned on the potential confounders age, sex, and ethnicity. Age is defined as the interval between sample extraction date and birthday and scaled to mean 0 and variance 1. If missing, months and days are set to the first day and month, respectively. Due to the ambiguous recording of ethnicity groups, samples are assigned to either African, Caucasian, or ‘Other’ ethnicity. Sex is modeled as a binary feature. Though deep sequencing has been performed, we use for this study only the consensus sequences derived using a 10% prevalence cutoff, which is commonly used in the research community [60]. The NGS reads were mapped with a customized version of MinVar [61].

Predicting if a polymorphism is under HLA pressure or not is confounded by the phylogenetic relatedness of the viral sequences. As proposed by [62], the phylogeny of the viral sequences of the subjects is incorporated as group-level effect (1|*subject*) into the model using the option cov_ranef = *list*(*subject* = *A*). Here, *A* denotes the computed covariance-matrix of the phylogenetic tree calculated using the vcv.phylo function from the ape package. A phylogenetic tree is constructed based on the nucleotide sequences of the p24 protein from the chronic_lowCD4 data set using the RAxML software (version 8.2.12) [63] under the GTRGAMMA model. Thus, the formula to compute the HLA model taking all HLA alleles *H* as potential covariates into the model has the following form:

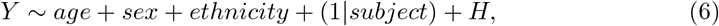

in contrast to the baseline model, which estimates the probability that the frequent site is not under HLA pressure:

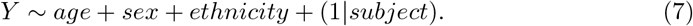

The logistic function is used as a link function. As described in previous sections, the horseshoe prior is used on all population-level effects [45]. Alleles in *H* are converted to four digit resolution. Alleles with alternative expression (suffix ‘L’, ‘S’, ‘C’, ‘A’, or ‘Q’) are treated separately from the normally expressed allele. The complete call to compute the per-site models using the brms package is provided in the code repository.

#### Calculation of adaptation score

We define the adaptation score, as proposed by [34], as:

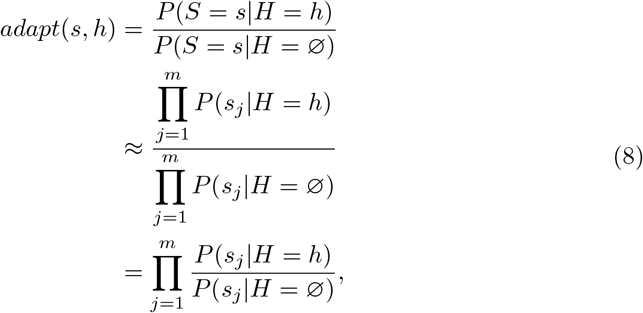

where the per-site likelihood *P*(*s_j_*|*H* = *h*) and *P*(*s_j_*|*H* = ∅) are defined by Eq.2 and Eq. 3, respectively. For better interpretation, we also transform the estimated adaptation *adapt*(*s*, *h*) using a sigmoidal function *g*(*x*) to a range of −1 to 1 [34]:

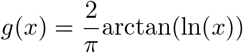

Thereby, a positive adaptation score denotes that the sequence has more likely occurred under HLA pressure than without, and vice versa.

#### Logo computation

The adaptation score can be decomposed into the likelihood ratios per frequent variant sites (see Eq. 8). Odds ratios above or below 1 indicate that either the polymorphism at the site *s_j_* is more likely to be under HLA pressure, or vice versa. We use this information to provide a visual logo depicting the amino acids that contributed most to the adaptation score. Therefore, only sites with odds ratios differing from 1 (and an offset of 0.01 to account for the variance) are considered. The contribution is scaled by the maximum contribution. In order to use the existing Weblogo 3.0 software to produce the logos [64], we create a pseudo-alignment with 100 sequences with length of the number of important sites. Each position in the alignment represents a polymorphism site. The sequences contain the polymorphism at this position with a frequency equal to the scaled contribution and a gap for the remaining sequences. Thus, the logo is a consensus logo for the pseudo-alignment.

### Data sets

We divide the newly sequenced study cohort based on a CD4^+^ T cell count cutoff of 500 cells/mm^3^ into a chronic_highCD4 data set and a chronic_lowCD4 count data set. High CD4^+^ T cell count indicates a stronger immune system. Since infection duration is not known for the patients, a high CD4^+^ T cell count might indicate that the patients have been infected for a shorter time (less chronic). Moreover, a virus is assumed to be less adapted to a host with a strong immune system compared to a host with a weak immune system. Thus, the adaptation model is only trained on the chronic_lowCD4 data set. In addition, we create an artificial data set (random) based on the chronic_lowCD4 data set, where the HLA alleles per HLA gene and haplotype have been randomized 100 times. HLA adaptation for this random data set is predicted with models based on the chronic_lowCD4 data set as well. For further validation of the adaptation model, we estimate HIV-1 adaptation of publicly available cohort of acutely-infected HIV-1 patients (n = 23) from the Los Alamos HIV sequence database (http://www.hiv.lanl.gov). The acute data set comprised the p24 sequence as well as the HLA I information of 23 patients with the following accession numbers GQ275453, GQ275750, GQ275852, GQ275894, KM192425, KM192440, KM192471, KM192536, KM192566, KM192640, KM192653, KM192674, KM192686, KM192702, KM192762, KM192844, KM192856, KM192870, KM192884, KM192912, KM192942, KM192970, KM192998. Since only the HLA I profile was available, we build an adaptation model based only on the HLA I profile for this purpose.

### Statistical analyses

We perform a one-sided Wilcoxon rank-sum test to compare the adaptation scores (i) between different data sets and (ii) with respect to different clinical characteristics. For settings, where the data is paired (random data set - same subjects, R5-FPR analysis - matched CD4 count, heterologous - autologous viruses), a one-sided Wilcoxon signed-rank test is performed. A significance threshold of 0.05 is set for all hypothesis tests.

### Data and code availability

The NGS sequences from the study cohort are available under the BioProject ID PRJNA810303 (reviewer link, see submission). The corresponding BioSample Accession IDs are SAMN26241863:26242168 and SAMN28728524:SAMN28728529. The generated consensus nucleotide sequences are provided on Zenodo at link 10.5281/zenodo.6797532. Due to privacy reasons, the HLA information cannot be published. Consequently, we cannot publish the trained models as the HLA information can be exposed thereby. A minimal data set including the estimated adaptation scores for all presented data sets is available on Zenodo at link 10.5281/zenodo.6797722. All code not compromising the privacy concerns, including the complete call to train and build the multinomial Bayesian generalized linear mixed models, is provided at GitHub at link https://github.com/annahake/HIVIA_TOOL.git to be used as template. To ensure reproducibility, we have used the workflow manager Snakemake 5.4.5 [65] and the Anaconda Software Distribution [66] for the training and prediction pipeline. We have used the R Language and Environment for Statistical Computing, Version 3.5.1 [55] for modeling and analyses.

## Results and Discussion

### Validation of the adaptation score

We trained our adaptation model on data from a cohort consisting of 274 chronically-infected, untreated, subtype C, HIV-1 patients, all having a CD4^+^ T cell count less than 500 cells/mm^3^ and on average a log viral load of 4.87 (‘chronic_lowCD4’ data set). In addition, 38 samples from the same study cohort with a CD4^+^ T cell count above 500 cells/mm^3^ (‘chronic_highCD4’ data set) were available. Apart from the CD4^+^ T cell count, the two data sets are comparable with regard to potential confounders and clinical variables (see Table 1).

**Table 1.** Summary statistics for the variables of interest for both data sets.

| variable | chronic_highCD4 | chronic_lowCD4 |
| --- | --- | --- |
| age | 35.24±10.03 | 35.38±9.99 |
| CD4<sup>+</sup> T cell count | 690.47±188.39 | 199.28±117.27 |
| VL | 4.22±0.73 | 4.87±0.85 |
| adapt(x) | 0.05±0.38 | 0.19±0.31 |
| sex:F | 0.58 | 0.61 |
| sex:M | 0.42 | 0.39 |
| ethnicity:AFRICAN | 0.79 | 0.64 |
| ethnicity:CAUCASIAN | 0.03 | 0 |
| ethnicity:OTHER | 0.18 | 0.36 |
| coreceptor:R5 | 0.82 | 0.71 |
| coreceptor:X4 | 0.18 | 0.29 |

Performing several runs of 10-fold cross-validation revealed that the predicted adaptation score is quite robust, changing with an average standard deviation around 0.1. Though there exists no ground truth for HLA adaptation, we set some requirements that a valid adaptation score should fulfill, which can be seen in the following.

#### Study cohort contains HLA adapted sequences

We assume that by construction the study cohort should harbor some HLA adapted sequences. 62% of the samples from the chronic_lowCD4 data set (n = 274) are estimated to be adapted (adaptation score >0.1), compared to 47% of the chronic_highCD4 data set (n = 38). The adaptation scores of the chronic_lowCD4 data set are taken from a 10-fold cross-validation, while the adaptation scores of the chronic_highCD4 data set are predicted using the full chronic_lowCD4 data set for training. Fig. 1 shows the distribution of the adaptation score in the chronic_lowCD4 data set and the chronic_lowCD4 data set. Statistically, HIV-1 isolates of patients with CD4^+^ T cell count below 500 (chronic_lowCD4 data set) are significantly more adapted than patients with higher CD4^+^ T cell count (one-sided, unpaired Wilcoxon rank-sum test, p-value = 1.97e-2). The comparison of the chronic_lowCD4 data set with the chronic_highCD4 data set is however not straightforward. On the one hand, the size of the chronic_highCD4 data set is quite small compared to the low CD4^+^ T cell. On the other hand, while we exclude the patients with the higher CD4^+^ T cell count from the training process as a precaution because they might be less chronic, this assumption does not have to be true and the samples cannot be treated to test the hypothesis that chronically-infected patients have more adapted viruses compared to patients with shorter infection duration. Last but not least, HLA-1 adapted viruses are assumed to escape the CTL response, resulting in fewer infected CD4^+^ T cells being killed. As a consequence, it is not necessarily the case that patients with higher CD4^+^ T cell count have less adapted viruses compared to patients with a lower CD4^+^ T cell count.

**Fig 1.**
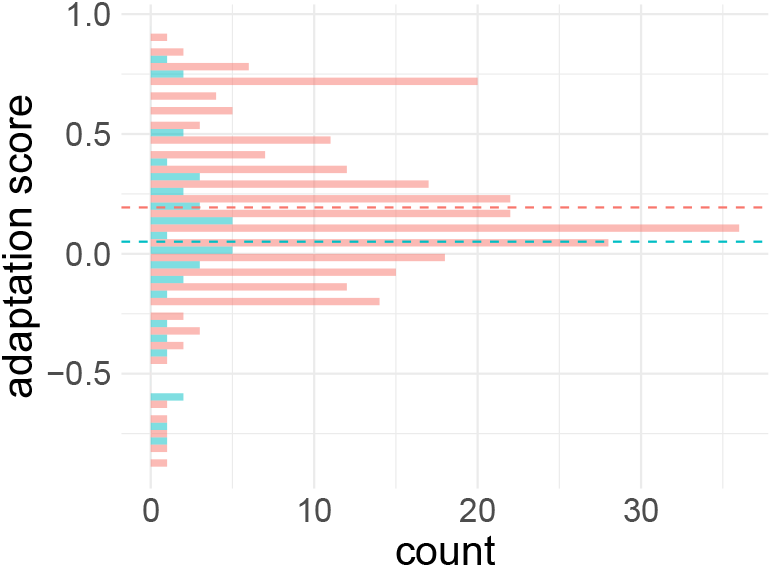
Histogram of the adaptation scores of the chronic_lowCD4 data set (red) and the chronic_highCD4 data set (turquoise). Dashed line represents the mean adaptation score per data set. The mean adaptation score is 0.19 for the chronic_lowCD4 data set and 0.05 for the chronic_highCD4 data set.

#### Random HLA profile leads to non-adaptedness

We expect that viruses in the study cohort are more adapted to the host’s HLA profile than to a random HLA profile. Therefore, we predicted the HLA adaptation of the viral sequences of the cohort to a random HLA profile (100 times). Adaptation scores in the random data set are averaged per patient over 100 draws. Only 10% of the random samples (n = 274) are predicted to be adapted. As expected, the adaptation of the same virus to a randomized HLA profile is significantly lower than to its host HLA profile (one-sided, paired Wilcoxon signed-rank test, p-value = 1.50e-44). Fig. 2 shows the distribution of the estimated adaptation scores for the random data set compared to the chronic_lowCD4 data set.

**Fig 2.**
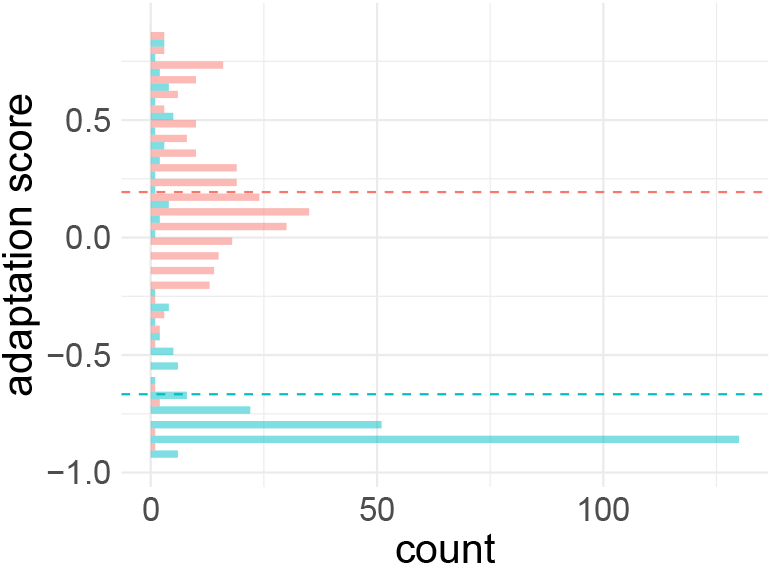
Histogram of the adaptation scores of the chronic_lowCD4 data set (red) and the the averaged adaptation scores of the random data set (turquoise). Dashed line represents the mean adaptation score per data set. The mean adaptation score is 0.19 for the chronic_lowCD4 data set and −0.67 for the random data set.

#### Autologous viruses more adapted than heterologous viruses

We observed that the adaptation score of the harbored virus to its host (autologous virus) is higher (p-value = 1.48e-31) in contrast to the adaptation of the other viruses in the cohort to the same HLA profile (heterologous virus). This meets our expectation, since we define the adaptation score to reflect how likely the virus acquired escape mutations specific to the host HLA profile. Fig. 3 shows the adaptation scores of the autologous virus and the averaged heterologous viruses for each subject (HLA profile).

**Fig 3.**
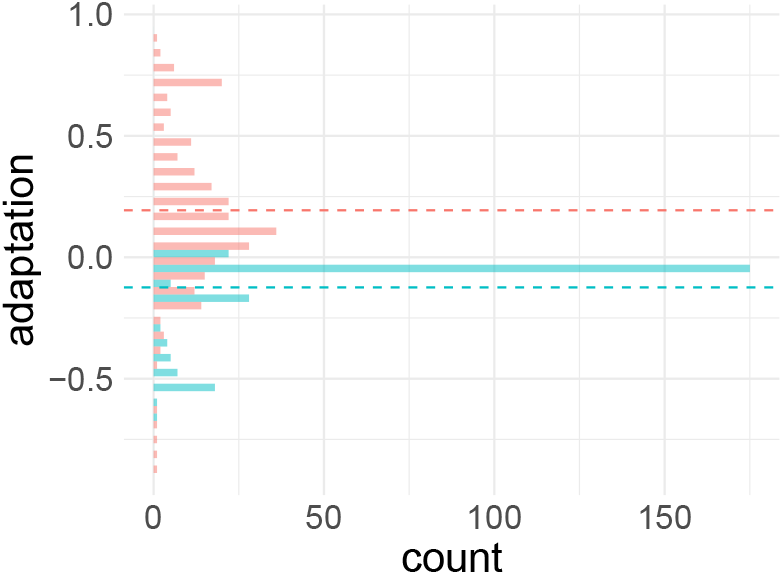
Histogram of estimated adapatation score for each HLA profile and autologous and heterologous viruses. Estimated adaptation scores for each HLA profile and its autologous virus (red) and heterologous viruses of the cohort (turquoise). The adaptation scores of the heterologous viruses are averaged. Dashed line represents the mean adaptation score per data set. The mean adaptation score for autologous viruses is 0.19 and −0.12 for heterologous viruses.

#### Viruses in acute phase less adapted than in chronic phase

We expect that viruses from acutely-infected HIV-1 patients should be less adapted than from chronically-infected HIV-1 patients due to the shorter exposure to the immune system. Fig. 4 shows a histogram of the estimated adaptation scores for the acute and the chronic data sets. Since only the HLA I profile was available for the acute data set, we built an adaptation model based only on the HLA I profile for this purpose. We observed that viral strains from acutely infected patients have significantly lower estimated adaptation scores compared to the chronically-infected HIV-1 patients from our cohort (one-sided, unpaired Wilcoxon rank-sum test, p-value = 4.17e-5). Note that viruses from acutely-infected patients might also carry HLA-related escape mutations due to transmission.

**Fig 4.**
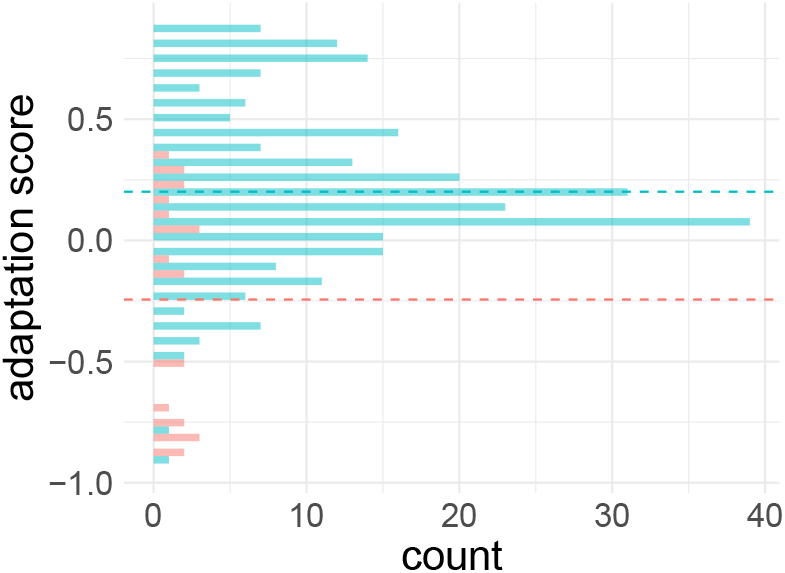
Comparison of HLA adaptation in acutely- and chronically-infected HIV-1 patients. Histogram of the estimated adaptation scores for the chronically-infected data set (turquoise) and the acutely infected data set (red). Dashed line represents the mean adaptation score per data set. The mean adaptation score is 0.20 for the chronic_lowCD4 data set based only on HLA I alleles and −0.24 for the acute data set, respectively.

### Validation of the per-site models

#### Non-informative per-site models have no influence on the adaptation score

In contrast to the overall adaptation score, it is possible to evaluate the performance of the per-site models. This is useful for the interpretation and validation of the model but irrelevant for the quality of the adaptation score. For each frequent site, we compute the likelihood ratio of a model that estimates the likelihood that the site is under HLA pressure (HLA model) and a hypothetical model that assumes no HLA pressure (baseline). Thereby, the estimated per-site adaptations are directly adjusted by a baseline model and calibrated among all sites. Thus, including sites which are not under HLA pressure will more likely contribute with a factor of 1 to the overall adaptation score and, consequently, have no influence. This allows to take all frequent sites into consideration without any preselection or apriori knowledge. Note, by definition of the adaptation score, the adaptation of each frequent site contributes with the same weight to the overall adaptation score. All per-site models reached the Gelman-Rubin convergence criteria by having an Rhat value less than or equal to 1.

#### Informative models learn HLA footprints

While it is not the focus of the study, we can identify sites with a likelihood ratio over 1, indicating a potential association between the frequent site and the HLA profile. In the study cohort, we identified 68 frequent sites in the p24 protein. Out of the corresponding 68 per-site HLA models, 21 had an averaged AUC under the precision-recall ROC curve higher than the averaged precision-recall baseline, where the precision-recall baseline is computed as the ratio of positive samples in the data set. Precision-recall was computed for each possible amino acid at a frequent site via 10-fold cross-validation. If models are evaluated by the performance to predict each frequent single amino acid polymorphism (SAP) separately, 52 models out of 210 perform better than the precision-recall baseline. Table 2 shows the top 10 polymorphisms with precision-recall AUC exceeding the baseline. Further analyzing the learned coefficients of the per-site models with high performance revealed that the models learned known footprints for subtype C such as the association between the T242N mutation and the HLA alleles HLA-B*57:01/02/03 or HLA-B*58:01 as well as the T186S escape mutation associated with HLA-B*81:01 [67–69].

**Table 2.** Top ten potential HLA-restricted sites and single amino acid polymorphisms (SAPs) with respect to precision-recall baseline performance. The performance of the HLA model at a specific site and for a specific SAP is computed as the AUC under the precision-recall curve (PRROC).

| site | SAP | PRROC | baseline |
| --- | --- | --- | --- |
| 242 | n | 0.81 | 0.12 |
| 186 | s | 0.45 | 0.04 |
| 163 | g | 0.26 | 0.06 |
| 309 | c | 0.20 | 0.01 |
| 146 | p | 0.38 | 0.24 |
| 357 | g | 0.65 | 0.53 |
| 242 | t | 0.98 | 0.87 |
| 146 | s | 0.21 | 0.11 |
| 312 | e | 0.39 | 0.30 |
| 230 | d | 0.14 | 0.05 |

#### Interpretable adaptation score by providing logos for each virus

For each frequent variant site, an odds ratio above or below 1 (with an offset of 0.1) indicates whether the amino acid at this site is more likely under HLA pressure or not. This information can be used to compute a logo revealing the amino acids that contributed the most to the adaptation score. This information helps the user to understand the results for different inputs. Fig. 5 shows the logo for the patient with the highest adaptation score in the cohort. The known HLA escape mutation T186S [70] has the highest contribution to the predicted adaptation score.

**Fig 5.**
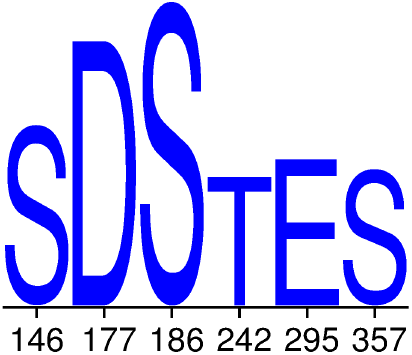
Logo for the patient with the highest adaptation score. The logo shows the viral polymorphisms that have the highest contribution to the adaptation score of this patient. Blue capital letters indicate adapted amino acids, while orange lowercase letters reflect non-adapted amino acids. The height of the letters reflects the contribution to the adaptation score and is scaled by the maximum contribution. The x-axis denotes the corresponding sites in the HXB2 virus.

### HLA adaptation associated with CD4^+^ T cell count but not viral load

We analyzed the estimated adaptation score with respect to viral load, CD4^+^ T cell count and coreceptor usage. On the one hand, we tested whether patients with adapted and non-adapted viruses differ in these variables, where adapted is defined as an adaptation score > 0.1 and non-adapted as an adaptation score < −0.1, based on the expected variance of 0.1 (see Fig. 6). On the other hand, we analyzed whether viruses of patients with different known levels of these variables differ in their adaptation (see Fig. 7).

**Fig 6.**
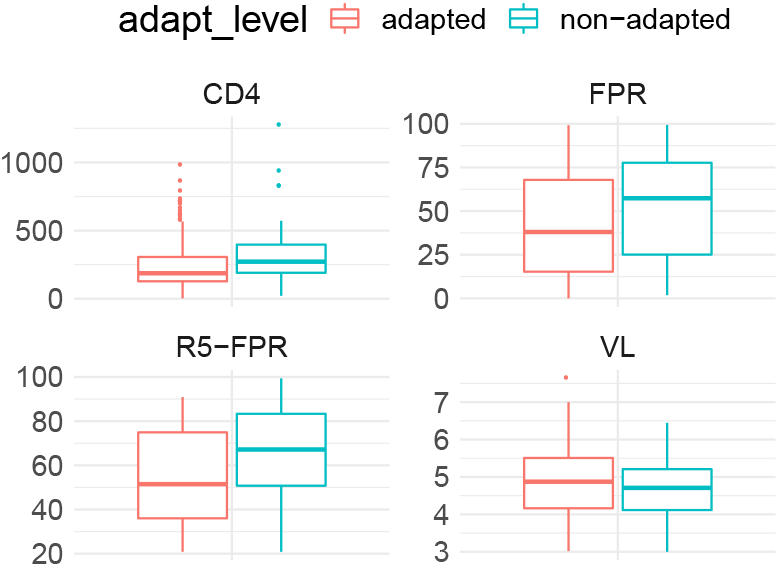
Difference in clinical variables based on HLA adaption. Measurement of CD4 ^+^ T cell (CD4), logarithmized viral load (VL), FPR, and the FPR of R5 viruses matched based on their CD4 count (R5-FPR) stratified among adapted (red) and non-adapted(turquoise) viruses.

**Fig 7.**
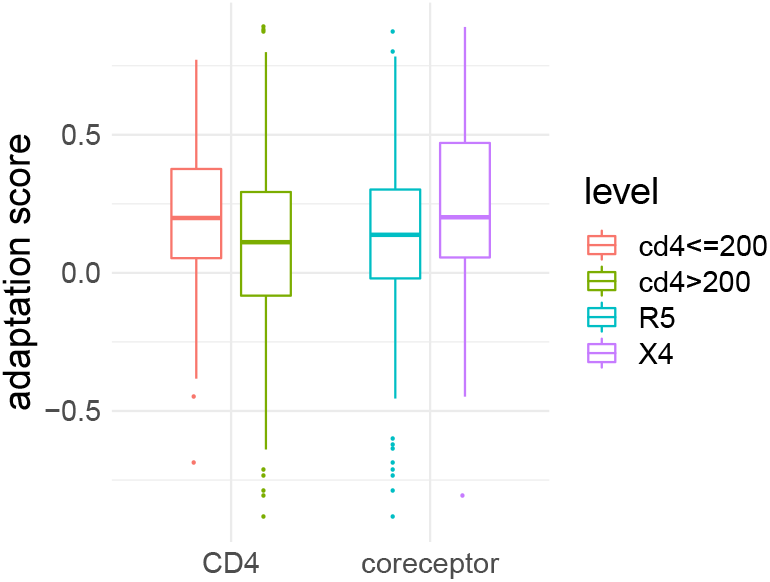
Adaptation score for different levels of CD4^+^ T cell count (CD4) and coreceptor usage (coreceptor)

Though HLA class-I restricted polymorphism are known to be predictive for viral load and CD4^+^ T cell count in general [26, 71], we observed only a correlation between the estimated adaptation scores (based on HLA I and HLA II alleles) and the CD4^+^ count (Pearson correlation coefficient −0.16, p-value=0.02) but not with viral load (0.04, p-value=0.88). Note, however, that the study cohort consists of rather chronically-infected patients at a later stage of infection, where other factors more likely affect fluctuations in the viral load than the HLA adaptation, and a difference between controllers and non-controllers, for example, is not expected to be seen as in the beginning of the infection. We also observed that adapted viruses do not have statistically significant higher viral loads than non-adapted viruses (one-sided, unpaired Wilcoxon rank-sum test, p-value = 1.86e-1), and that patients with low viral load have not less adapted viruses (one-sided, unpaired Wilcoxon rank-sum test, p-value=8.54e-2). In addition to the significant correlation between the CD4^+^ T cell count and adaptation score, we observed that patients with AIDS (CD4^+^ T cell count < 200) have more adapted viruses than patients with higher CD4^+^ T cell counts (one-sided, unpaired, Wilcoxon rank-sum test, p-value = 3.20e-3). CD4^+^ T cell count was also lower in patients with adapted viruses compared to non-adapted (Wilcoxon rank-sum test, p-value = 1.27e-3).

### Adaptation associated with coreceptor usage

Using our adaptation score, we investigated the relationship between HLA adaptation and coreceptor usage. More precisely, we analyzed the hypothesis that high HLA adaptation might trigger the coreceptor switch in a similar way as a weak immune system (measured by a low number of CD4^+^ T cell counts). Coreceptor usage was determined with the false positive rate (FPR) of the coreceptor prediction tool geno2pheno[coreceptor] [17]. The provided FPR corresponds to the confidence with which the sequence is classified as X4-capable. The higher the FPR, the more likely the sequence is not X4-capable, but R5-capable. Table 3 shows the average adaptation scores stratified for coreceptor usage. We observed a negative correlation between estimated adaptation score and corresponding FPR (Pearson correlation coefficient of −0.15, p-value = 0.03). This means that the more adapted the virus, the higher the likelihood that the virus is classified as X4-capable. This was further confirmed by the observation that X4-capable viruses are more adapted compared to R5 viruses (Wilcoxon rank-sum test, p-value = 1.34e-2, see Fig. 7) and that, in general, adapted viruses have a lower FPR (rather X4 variants) compared to non-adapted viruses (Wilcoxon rank-sum test, p-value = 6.76e-3, see Fig. 6). Note, since the variants are already determined as *X*4-capable, it is impossible to show if the emergence of X4-variants is driven by HLA adaptation. This analysis would require longitudinal data where the emergence of the coreceptor switch is captured. To rule out the possibility that higher adaptation of the X4 variants occurs due to longer exposure to the host immune system in contrast to R5 variants, the exact duration of infection is required. However, we observed that even among all R5 viruses, higher adaptation is associated with lower FPR, indicating that more adapted R5 samples might be closer to the coreceptor switch compared to non-adapted samples (one-sided paired Wilcoxon signed-rank test, p-value = 2.21e-2). Since the CD4^+^ T cell count is a major confounder for the coreceptor usage, we have matched for this test adapted and non-adapted R5 samples with similar CD4^+^ count (± 50 cells/mm^3^). Note, high adaptation of an R5 variant in a chronically-infected patient can also occur due to the long exposure to the immune system, since a coreceptor switch is only observed in 50% of the patients.

**Table 3.** Averaged and maximum adaptation score stratified on the coreceptor usage and two data sets.

| data set | coreceptor | adapt(x) | max(adap(x)) |
| --- | --- | --- | --- |
| chronic_highCD4 | R5 | 0.03±0.40 | 0.80 |
| chronic_highCD4 | X4 | 0.14±0.28 | 0.51 |
| chronic_lowCD4 | R5 | 0.17±0.30 | 0.87 |
| chronic_lowCD4 | X4 | 0.25±0.33 | 0.89 |

## Conclusion

Here, we introduced a novel computational approach to jointly estimate HLA I and HLA II adaptation of HIV-1 using Bayesian generalized linear mixed models. In addition, we presented a new study cohort of 312 treatment-naive, subtype C, chronically-infected HIV-1 patients from South Africa, where we sequenced the viral *gag* (and *env*) protein with corresponding HLA class I and II alleles for the training of our models. Apart from validating that our adaptation score inherited appropriate characteristics, we showed that the models underlying the adaptation score are biologically meaningful by learning well-known HLA-restricted polymorphisms. Using our approach and the data, we investigated the relationship between HLA adaptation and coreceptor usage of HIV-1, which had been unexplored up to now. We observed that X4-capable viruses are more adapted compared to R5-capable viruses (Wilcoxon rank-sum test, p-value =1.34e-2). Moreover, even among all R5-capable viruses, higher adaptation is associated with lower FPR, indicating that more adapted R5 variants might be closer to the coreceptor switch compared to non-adapted variants (Wilcoxon signed-rank test, p-value = 2.21e-2). Thus, HLA adaptation might be another factor that should be considered prior to the administration of CCR5 coreceptor antagonists. It might also be useful in predicting how imminent the coreceptor switch is.

In general, the estimated adaptation score allows to measure and understand HIV-1’s adaptation to the immune system. The adaptation score can be used to guide the design of suitable immunogens as vaccine targets by selecting sites that are more likely to be non-adapted to the immune system. Since the approach itself is not HIV-1 specific, the presented method can be also applied to study any virus-host adaptation. We encourage the usage of Bayesian GLMMs for modeling virus-host adaptation due to their ability to adjust for phylogenetic dependencies in the data and to handle highly overparameterized settings within the model. In light of current and potential future viral threats to mankind, such as SARS-CoV-2 or Ebola or MERS-CoV, this flexible, data non-intensive method can be useful to reveal and analyze virus-host dynamics of new viruses where little data is available.

Future studies of the study cohort are required to further evaluate how the adaptation score is coherent with CTL escape experiments. While the study cohort was appropriate to learn HLA adaptation, it only allows to study the coreceptor switch and the role of HLA adaptation on it from a retrospective angle. Moreover, the number of CXCR4-using variants with intermediate to high CD4^+^ T cell count was very low. Consequently, a study cohort with longitudinal data on coreceptor usage and intermediate to high CD4^+^ T cell count would give additional insights. This would not only allow to investigate if HLA II adaptation occurs prior to the coreceptor switch, but also if the degree of adaptation is associated with the time until the coreceptor switch occurs.

Note that the presented adaptation score here is a simple approach that can be easily optimized and extended in different ways. Given the available data, we restricted the analysis to subtype C infected patients and the p24 protein. However, our approach can also be applied to other subtypes and/or combined over different viral proteins. Further, we used the viral consensus sequences instead of the NGS sequences, since we aimed at predicting the adaptation per sequence. Still, the within-subject relatedness of different reads per virus could be easily incorporated into the Bayesian models. A larger data set might improve the current adaptation score by better representing the population with respect to the HLA repertoire and potential frequent variant sites, resulting in more informative per-site models.

Furthermore, it is possible to make the proposed models more complex by incorporating more dependency structures such as HLA linkage, or by relaxing the assumption of independence among all sites to capture compensatory mutations. Another assumption is that each frequent variant site has the same probability to be under HLA pressure. Prior knowledge about common HLA epitopes can be added to the model by weighting the per-site likelihood odds ratios accordingly. However, if a site is more likely to be under HLA pressure, given by the underlying data, by construction of the adaptation score, the likelihood odds ratio should contribute with a higher factor to the overall adaptation score.

Decomposing the adaptation score based on the potential adaptation of each frequent variant site is very advantageous with respect to model explainability and to settings, where little prior information exists. However, it requires the computation of two models per frequent variant sites, leading to a high number of models. Currently, the per-site models are not optimized with regard to parameter and predictor selection. To avoid overfitting and p-value based selection, we forced each model to capture our prior beliefs that the model should be unbiased with regard to sex, age, ethnicity and not be hampered by phylogenetic relatedness. While we ensured that the models are all converging according to the Rubin-Gelman criterion, we do not perform visual checks of the Bayesian GLMMs, such as prior and posterior predictive checks. Though this is a standard procedure for Bayesian GLMMs, it was not feasible in our case due to the high number of models. In our case, this was also not mandatory. Setting the horseshoe prior for the beta coefficients was based on our apriori knowledge that only few HLA alleles and clinical variables should have influence on a site. Setting potential non-optimal parameters might lead to non-informative per-site models. While we might lose some potential information for these sites, the quality of the overall adaptation score remains guaranteed by calibrating the per-site models with a baseline model. For computational reasons, it might be also beneficial to reduce the computation of the adaptation score based on only the informative per-site models.

## References

1. on HIV/AIDS JUNP, et al. Fast-track: ending the AIDS epidemic by 2030. Geneva: UNAIDS. 2014;.

2. Lusso P. HIV and the chemokine system: 10 years later. The EMBO Journal. 2006;25(3):447–456. doi:10.1038/sj.emboj.7600947.

3. Berger EA, Doms R, Fenyö EM, Korber B, Littman D, Moore J, et al. A new classification for HIV-1. Nature. 1998;391(6664):240–240.

4. Moore JP, Kitchen SG, Pugach P, Zack JA. The CCR5 and CXCR4 Coreceptors—Central to Understanding the Transmission and Pathogenesis of Human Immunodeficiency Virus Type 1 Infection. AIDS Research and Human Retroviruses. 2004;20(1):111–126. doi:10.1089/088922204322749567.

5. Keele BF, Giorgi EE, Salazar-Gonzalez JF, Decker JM, Pham KT, Salazar MG, et al. Identification and characterization of transmitted and early founder virus envelopes in primary HIV-1 infection. Proceedings of the National Academy of Sciences. 2008;105(21):7552–7557. doi:10.1073/pnas.0802203105.

6. Connor RI, Sheridan KE, Ceradini D, Choe S, Landau NR. Change in Coreceptor Use Correlates with Disease Progression in HIV-1–Infected Individuals. Journal of Experimental Medicine. 1997;185(4):621–628. doi:10.1084/jem.185.4.621.

7. Regoes RR, Bonhoeffer S. The HIV coreceptor switch: a population dynamical perspective. Trends in microbiology. 2005;13(6):269–277.

8. Schuitemaker H, Koot M, Kootstra NA, Dercksen MW, de Goede RE, van Steenwijk RP, et al. Biological phenotype of human immunodeficiency virus type 1 clones at different stages of infection: progression of disease is associated with a shift from monocytotropic to T-cell-tropic virus population. Journal of Virology. 1992;66(3):1354–1360.

9. Tscherning C, Alaeus A, Fredriksson R, Åsa Björndal, Deng H, Littman DR, et al. Differences in Chemokine Coreceptor Usage between Genetic Subtypes of HIV-1. Virology. 1998;241(2):181–188. doi:https://doi.org/10.1006/viro.1997.8980.

10. Connell BJ, Michler K, Capovilla A, Venter WDF, Stevens WS, Papathanasopoulos MA. Emergence of X4 usage among HIV-1 subtype C: evidence for an evolving epidemic in South Africa. AIDS. 2008;22:896–899.

11. Whitcomb JM, Huang W, Fransen S, Limoli K, Toma J, Wrin T, et al. Development and Characterization of a Novel Single-Cycle Recombinant-Virus Assay To Determine Human Immunodeficiency Virus Type 1 Coreceptor Tropism. Antimicrobial Agents and Chemotherapy. 2006;51(2):566–575. doi:10.1128/aac.00853-06.

12. Low AJ, McGovern RA, Harrigan PR. Trofile HIV co-receptor usage assay. Expert opinion on medical diagnostics. 2009;3(2):181–191.

13. Reeves J, Coakley E, Petropoulos C, Whitcomb J. An enhanced sensitivity Trofile HIV coreceptor tropism assay for selecting patients for therapy with entry inhibitors targeting CCR5: a review of analytical and clinical studies. J Viral Entry. 2009;3(3):94–102.

14. Gonzalez-Serna A, Leal M, Genebat M, Abad MA, Garcia-Perganeda A, Ferrando-Martinez S, et al. TROCAI (Tropism Coreceptor Assay Information): a New Phenotypic Tropism Test and Its Correlation with Trofile Enhanced Sensitivity and Genotypic Approaches. Journal of Clinical Microbiology. 2010;48(12):4453–4458. doi:10.1128/JCM.00953-10.

15. Fouchier R, Groenink M, Kootstra NA, Tersmette M, Huisman H, Miedema F, et al. Phenotype-associated sequence variation in the third variable domain of the human immunodeficiency virus type 1 gp120 molecule. Journal of virology. 1992;66(5):3183–3187.

16. Jensen MA, Li FS, van’t Wout AB, Nickle DC, Shriner D, He HX, et al. Improved coreceptor usage prediction and genotypic monitoring of R5-to-X4 transition by motif analysis of human immunodeficiency virus type 1 env V3 loop sequences. Journal of virology. 2003;77(24):13376–13388.

17. Lengauer T, Sander O, Sierra S, Thielen A, Kaiser R. Bioinformatics prediction of HIV coreceptor usage. Nature biotechnology. 2007;25(12):1407–1410.

18. Pfeifer N, Lengauer T. Improving HIV coreceptor usage prediction in the clinic using hints from next-generation sequencing data. Bioinformatics. 2012;28(18):i589–i595. doi:10.1093/bioinformatics/bts373.

19. Cashin K, Gray LR, Jakobsen MR, Sterjovski J, Churchill MJ, Gorry PR. CoRSeqV3-C: a novel HIV-1 subtype C specific V3 sequence based coreceptor usage prediction algorithm. Retrovirology. 2013;10(1):24. doi:10.1186/1742-4690-10-24.

20. Sander O, Sing T, Sommer I, Low AJ, Cheung PK, Harrigan PR, et al. Structural descriptors of gp120 V3 loop for the prediction of HIV-1 coreceptor usage. PLoS computational biology. 2007;3(3).

21. Sing T, Low AJ, Beerenwinkel N, Sander O, Cheung PK, Domingues FS, et al. Predicting HIV coreceptor usage on the basis of genetic and clinical covariates. Antiviral therapy. 2007;12(7):1097.

22. Swenson LC, Mo T, Dong WW, Zhong X, Woods CK, Jensen MA, et al. Deep sequencing to infer HIV-1 co-receptor usage: application to three clinical trials of maraviroc in treatment-experienced patients. Journal of Infectious Diseases. 2011;203(2):237–245.

23. Pollakis G, Kang S, Kliphuis A, Chalaby MI, Goudsmit J, Paxton WA. N-linked glycosylation of the HIV type-1 gp120 envelope glycoprotein as a major determinant of CCR5 and CXCR4 coreceptor utilization. Journal of Biological Chemistry. 2001;276(16):13433–13441.

24. Phillips RE, Rowland-Jones S, Nixon DF, Gotch FM, Edwards JP, Ogunlesi AO, et al. Human immunodeficiency virus genetic variation that can escape cytotoxic T cell recognition. Nature. 1991;354(6353):453–459.

25. Goulder PJR, Watkins DI. HIV and SIV CTL escape: implications for vaccine design. Nature Reviews Immunology. 2004;4(8):630–640. doi:10.1038/nri1417.

26. The International HIV Controllers Study. The Major Genetic Determinants of HIV-1 Control Affect HLA Class I Peptide Presentation. Science. 2010;330(6010):1551–1557. doi:10.1126/science.1195271.

27. Rousseau CM, Daniels MG, Carlson JM, Kadie C, Crawford H, Prendergast A, et al. HLA Class I-Driven Evolution of Human Immunodeficiency Virus Type 1 Subtype C Proteome: Immune Escape and Viral Load. Journal of Virology. 2008;82(13):6434–6446. doi:10.1128/jvi.02455-07.

28. Carlson JM, Brumme ZL, Rousseau CM, Brumme CJ, Matthews P, Kadie C, et al. Phylogenetic Dependency Networks: Inferring Patterns of CTL Escape and Codon Covariation in HIV-1 Gag. PLoS Computational Biology. 2008;4(11):e1000225. doi:10.1371/journal.pcbi.1000225.

29. Moore C, John M, James I, Christiansen F, Witt C, Mallal S. Evidence of HIV-1 adaptation to HLA-restricted immune responses at a population level. Science (New York, NY). 2002;296:1439–43.

30. Fellay J, Shianna KV, Ge D, Colombo S, Ledergerber B, Weale M, et al. A Whole-Genome Association Study of Major Determinants for Host Control of HIV-1. Science. 2007;317(5840):944–947. doi:10.1126/science.1143767.

31. Carlson JM, Brumme CJ, Martin E, Listgarten J, Brockman MA, Le AQ, et al. Correlates of protective cellular immunity revealed by analysis of population-level immune escape pathways in HIV-1. Journal of virology. 2012;86(24):13202–13216. doi:10.1128/JVI.01998-12.

32. Migueles SA, Sabbaghian MS, Shupert WL, Bettinotti MP, Marincola FM, Martino L, et al. HLA B*5701 is highly associated with restriction of virus replication in a subgroup of HIV-infected long term nonprogressors. Proceedings of the National Academy of Sciences. 2000;97(6):2709–2714. doi:10.1073/pnas.050567397.

33. Altfeld M, Addo MM, Rosenberg ES, Hecht FM, Lee PK, Vogel M, et al. Influence of HLA-B57 on clinical presentation and viral control during acute HIV-1 infection. AIDS. 2003;17(18).

34. Carlson JM, Du VY, Pfeifer N, Bansal A, Tan VYF, Power K, et al. Impact of pre-adapted HIV transmission. Nature Medicine. 2016;22(6):606–613. doi:10.1038/nm.4100.

35. Erdmann N, Du VY, Carlson J, Schaefer M, Jureka A, Sterrett S, et al. HLA Class-II Associated HIV Polymorphisms Predict Escape from CD4+ T Cell Responses. PLOS Pathogens. 2015;11(8):e1005111. doi:10.1371/journal.ppat.1005111.

36. Bhattacharya T, Daniels M, Heckerman D, Foley B, Frahm N, Kadie C, et al. Founder Effects in the Assessment of HIV Polymorphisms and HLA Allele Associations. Science. 2007;315(5818):1583–1586. doi:10.1126/science.1131528.

37. Troyer RM, McNevin J, Liu Y, Zhang SC, Krizan RW, Abraha A, et al. Variable fitness impact of HIV-1 escape mutations to cytotoxic T lymphocyte (CTL) response. PLoS pathogens. 2009;5(4).

38. Payne R, Branch S, Kløverpris H, Matthews P, Koofhethile C, Strong T, et al. Differential escape patterns within the dominant HLA-B* 57: 03-restricted HIV Gag epitope reflect distinct clade-specific functional constraints. Journal of virology. 2014;88(9):4668–4678.

39. Prince JL, Claiborne DT, Carlson JM, Schaefer M, Yu T, Lahki S, et al. Role of transmitted Gag CTL polymorphisms in defining replicative capacity and early HIV-1 pathogenesis. PLoS pathogens. 2012;8(11).

40. Kiepiela P, Ngumbela K, Thobakgale C, Ramduth D, Honeyborne I, Moodley E, et al. CD8+ T-cell responses to different HIV proteins have discordant associations with viral load. Nature medicine. 2007;13(1):46–53.

41. Edwards BH, Bansal A, Sabbaj S, Bakari J, Mulligan MJ, Goepfert PA. Magnitude of Functional CD8+ T-Cell Responses to the Gag Protein of Human Immunodeficiency Virus Type 1 Correlates Inversely with Viral Load in Plasma. Journal of Virology. 2002;76(5):2298–2305. doi:10.1128/jvi.76.5.2298-2305.2002.

42. Geldmacher C, Currier JR, Herrmann E, Haule A, Kuta E, McCutchan F, et al. CD8 T-Cell Recognition of Multiple Epitopes within Specific Gag Regions Is Associated with Maintenance of a Low Steady-State Viremia in Human Immunodeficiency Virus Type 1-Seropositive Patients. Journal of Virology. 2006;81(5):2440–2448. doi:10.1128/jvi.01847-06.

43. Zuniga R, Lucchetti A, Galvan P, Sanchez S, Sanchez C, Hernandez A, et al. Relative Dominance of Gag p24-Specific Cytotoxic T Lymphocytes Is Associated with Human Immunodeficiency Virus Control. Journal of Virology. 2006;80(6):3122–3125. doi:10.1128/jvi.80.6.3122-3125.2006.

44. Kløverpris HN, Leslie A, Goulder P. Role of HLA Adaptation in HIV Evolution. Frontiers in Immunology. 2016;6. doi:10.3389/fimmu.2015.00665.

45. Carvalho CM, Polson NG, Scott JG. Handling sparsity via the horseshoe. Journal of Machine Learning Research W&CP. 2009;.

46. Struck D, Lawyer G, Ternes AM, Schmit JC, Bercoff DP. COMET: adaptive context-based modeling for ultrafast HIV-1 subtype identification. Nucleic acids research. 2014;42(18):e144. doi:10.1093/nar/gku739.

47. Porter D, Daeumer M, Thielen A, Chang S, Martin R, Cohen C, et al. Emergent HIV-1 Drug Resistance Mutations Were Not Present at Low-Frequency at Baseline in Non-Nucleoside Reverse Transcriptase Inhibitor-Treated Subjects in the STaR Study. Viruses. 2015;7(12):6360–6370. doi:10.3390/v7122943.

48. Vandekerckhove L, Wensing A, Kaiser R, Brun-Vézinet F, Clotet B, De Luca A, et al. European guidelines on the clinical management of HIV-1 tropism testing. The Lancet infectious diseases. 2011;11(5):394–407.

49. Katoh K, Standley DM. MAFFT multiple sequence alignment software version 7: improvements in performance and usability. Molecular biology and evolution. 2013;30(4):772–80. doi:10.1093/molbev/mst010.

50. Nelder JA, Wedderburn RW. Generalized linear models. Journal of the Royal Statistical Society: Series A (General). 1972;135(3):370–384.

51. McCullagh P. Generalized linear models. Routledge; 2018.

52. Agresti A. An Introduction to Categorical Data Analysis. John Wiley & Sons, Inc.; 2007. Available from: https://doi.org/10.1002/0470114754.

53. Fong Y, Rue H, Wakefield J. Bayesian inference for generalized linear mixed models. Biostatistics. 2009;11(3):397–412. doi:10.1093/biostatistics/kxp053.

54. Fox J, Weisberg S. An R Companion to Applied Regression. 3rd ed. Thousand Oaks CA: Sage; 2019. Available from: http://tinyurl.com/carbook.

55. R Core Team. R: A Language and Environment for Statistical Computing; 2017. Available from: https://www.R-project.org/.

56. Bürkner PC. brms: An R Package for Bayesian Multilevel Models Using Stan. Journal of Statistical Software, Articles. 2017;80(1):1–28. doi:10.18637/jss.v080.i01.

57. Carpenter B, Gelman A, Hoffman M, Lee D, Goodrich B, Betancourt M, et al. Stan: A Probabilistic Programming Language. Journal of Statistical Software, Articles. 2017;76(1):1–32. doi:10.18637/jss.v076.i01.

58. Duane S, Kennedy AD, Pendleton BJ, Roweth D. Hybrid Monte Carlo. Physics Letters B. 1987;195(2):216–222. doi:10.1016/0370-2693(87)91197-x.

59. Homan MD, Gelman A. The No-U-Turn Sampler: Adaptively Setting Path Lengths in Hamiltonian Monte Carlo. J Mach Learn Res. 2014;15(1):1593–1623.

60. Dóring M, Büch J, Friedrich G, Pironti A, Kalaghatgi P, Knops E, et al. geno2pheno [ngs-freq]: a genotypic interpretation system for identifying viral drug resistance using next-generation sequencing data. Nucleic Acids Research. 2018;.

61. Huber M, Metzner KJ, Geissberger FD, Shah C, Leemann C, Klimkait T, et al. MinVar: A rapid and versatile tool for HIV-1 drug resistance genotyping by deep sequencing. Journal of Virological Methods. 2017;240:7–13. doi:https://doi.org/10.1016/j.jviromet.2016.11.008.

62. Hadfield JD, Nakagawa S. General quantitative genetic methods for comparative biology: phylogenies, taxonomies and multi-trait models for continuous and categorical characters. Journal of Evolutionary Biology. 2010;23(3):494–508. doi:10.1111/j.1420-9101.2009.01915.x.

63. Stamatakis A. RAxML version 8: a tool for phylogenetic analysis and post-analysis of large phylogenies. Bioinformatics. 2014;30(9):1312–1313. doi:10.1093/bioinformatics/btu033.

64. Schneider TD, Stephens RM. Sequence Logos: A New Way to Display Consensus Sequences. Nucleic Acids Res. 1990;18:6097–6100.

65. Mólder F, Jablonski KP, Letcher B, Hall MB, Tomkins-Tinch CH, Sochat VV, et al. Sustainable data analysis with Snakemake. F1000Research. 2021;10.

66. Anaconda Software Distribution; 2020. Available from: https://docs.anaconda.com/.

67. Wright JK, Naidoo VL, Brumme ZL, Prince JL, Claiborne DT, Goulder PJR, et al. Impact of HLA-B*81-Associated Mutations in HIV-1 Gag on Viral Replication Capacity. Journal of Virology. 2012;86(6):3193–3199. doi:10.1128/jvi.06682-11.

68. Leslie A, Matthews PC, Listgarten J, Carlson JM, Kadie C, Ndung’u T, et al. Additive Contribution of HLA Class I Alleles in the Immune Control of HIV-1 Infection. Journal of Virology. 2010;84(19):9879–9888. doi:10.1128/jvi.00320-10.

69. Kiepiela P, Leslie AJ, Honeyborne I, Ramduth D, Thobakgale C, Chetty S, et al. Dominant influence of HLA-B in mediating the potential co-evolution of HIV and HLA. Nature. 2004;432(7018):769–775.

70. Kløverpris HN, Leslie A, Goulder P. Role of HLA Adaptation in HIV Evolution. Frontiers in Immunology. 2016;6. doi:10.3389/fimmu.2015.00665.

71. Carrington M, O’Brien SJ. The Influence of HLA Genotype on AIDS. Annual Review of Medicine. 2003;54(1):535–551. doi:10.1146/annurev.med.54.101601.152346.

